# Alternative splicing in neuroblastoma generates RNA-fusion transcripts and is associated with vulnerability to spliceosome inhibitors

**DOI:** 10.1101/851238

**Authors:** Yao Shi, Vilma Rraklli, Eva Maxymovitz, Shuijie Li, Isabelle Westerlund, Oscar C. Bedoya-Reina, Juan Yuan, Petra Bullova, C. Christofer Juhlin, Adam Stenman, Catharina Larsson, Per Kogner, Maureen J. O’Sullivan, Susanne Schlisio, Johan Holmberg

## Abstract

The paucity of recurrent mutations has hampered efforts to understand and treat neuroblastoma. Alternative splicing and splicing-dependent RNA-fusions represent mechanisms able to increase the gene product repertoire but their role in neuroblastoma remains largely unexplored. Through analysis of RNA-sequenced neuroblastoma we here show that elevated expression of splicing factors is a strong predictor of poor clinical outcome. Furthermore, we identified >900 primarily intrachromosomal fusions containing canonical splicing sites. Fusions included transcripts from well-known oncogenes, were enriched for proximal genes and in chromosomal regions commonly gained or lost in neuroblastoma. As a proof-of-principle that these fusions can generate altered gene products, we characterized a *ZNF451*-*BAG2* fusion, generating a truncated BAG2-protein which inhibited retinoic acid-induced differentiation. Spliceosome inhibition impeded neuroblastoma fusion expression, induced apoptosis and inhibited xenograft tumor growth. Our findings elucidate a splicing-dependent mechanism producing altered gene products in neuroblastoma and suggest that the spliceosome is a tractable therapeutical target.

## Introduction

Neuroblastoma arises within the sympathetic nervous system and is the most frequent extracranial solid childhood cancer, exhibiting a high degree of clinical heterogeneity ranging from spontaneous regression to fatal progression (1). Despite intense sequencing efforts, few recurrently mutated genes have been identified in neuroblastoma (2, 3). Instead, high-risk neuroblastoma are characterized by large-scale chromosomal rearrangements such as chromothripsis and loss or gain of chromosomal regions (e.g. loss of 1p36 and 11q or gain of 17q and 2p) with or without *MYCN* amplification (*1, 2*). In eukaryotic cells, pre-mRNA splicing is essential for intron removal and to increase the number of possible gene products. If not properly regulated, this process can lead to mis-splicing, generating altered and potentially oncogenic proteins. A hyperactivated spliceosome can also provide an avenue of intervention, e.g. in *MYC*-driven breast cancer, where an increase in precursor mRNA synthesis results in an increased burden on the splicing machinery and consequently a vulnerability to spliceosome inhibition (*4*). In addition, several cancers harbor monoallelic mutations of spliceosome factors (*5*). Given the lack of recurring mutations and the prominent role of *MYCN* for high-risk NB, understanding a possible role for altered splicing dynamics in high-risk NB could provide novel insights on how to target this disease. Notably, in a mouse-model of neuroblastoma, intronic splicing motifs was shown to represent hotspots for recurring mutations (*6*). A potential consequence of aberrant splicing is generation of fusion transcripts through *cis*-splicing of adjacent genes (*7, 8*). Such fusions differ from those generated by chromosomal translocations (e.g. BCR-ABL in chronic myelogenous leukemia (*9*)). *Cis*-spliced fusions represent an alternative mechanism, distinct from amino acid changing mutations, able to generate altered gene products with a potential to promote tumors but also serve as diagnostic markers or novel drug targets (*7, 10*). Besides a fusion resulting from small interstitial genomic deletions at 11q generating either a *MLL-FOXR1* or a *PAFAH1B2*-*FOXR2* fusion (*11*) no intra-chromosomal chimeric transcripts have been described in neuroblastoma. However, they have been shown to be present in other tumor types as well as in non-transformed tissues and be promoted by different types of cellular stress such as infections or mutations (*12–16*).

In order to explore whether neuroblastoma tumors harbor previously undetected gene fusions, we analyzed a cohort of 172 RNA-sequenced neuroblastoma tumors. We identified several fusion transcripts, of which a significant proportion exhibited a distinct genomic distribution according to tumor risk. As a proof of principle, that fusions can generate novel gene products with alternative properties, we cloned and characterized the *ZNF451-BAG2* fusion. The generated protein exhibited distinct protein-protein binding properties compared to wild-type BAG2 and impeded retinoic acid induced differentiation. This reveals how a fusion gene product can influence neuroblastoma response to a drug commonly used in the treatment of high-risk patients (*17*). Identified fusions were predominantly generated by genes in close proximity and flanked by canonical splicing donors and acceptors. This pattern was distinct to fusions unique for neuroblastoma, whereas fusions we identified in normal adrenal gland or in other tumors lacked such a pattern. Furthermore, a subset of identified NB specific fusions was hypersensitive to pharmacologic spliceosome inhibition in comparison to their wild type cognates. High expression levels of spliceosome factors were strongly associated with high-risk disease and spliceosome inhibition also selectively promoted apoptosis in neuroblastoma cells *in vitro* and reduced tumor growth in mouse xenografts.

## Results

### Fusion transcripts are a common feature of neuroblastoma

To reveal novel gene fusions in neuroblastoma we analyzed a data set (National Cancer Institute TARGET, dbGap Study Accession: phs000218.v16.p6) comprising 172 paired-end RNA sequenced neuroblastoma tumors (referred to as “NB172”), out of which 139 were diagnosed as high-risk, 19 as intermediate-risk and 14 as low-risk according to the Children’s Oncology Groups staging (COG), (Supplementary Table 1). We applied the fusion detection tool FusionCatcher (*18*) and identified chimeric transcripts in 163 out of 172 cases with an average of 31 distinct fusion transcripts per tumor (Supplementary Table 2). Short homologous sequences (SHS) have been suggested to serve as templates for reverse transcriptase dependent false positive chimeras/fusions (*19*). In order to avoid potential false positive fusions, we removed any fusion that incorporated genes with SHSs of five or more nucleotides, which reduced the number of identified fusions from 1073 to 924. Structural consequences of the fusions ranged from truncated proteins through bona fide fusion proteins to deletion of genes. The majority of fusions (786/924) revealed by our analysis were intra-chromosomal fusion transcripts (Supplementary Table 2-3) many of which consisted of adjacent genes. Importantly, 114 fusions occurred at a frequency of 5% or more (Top 25 in Fig. 1A, all >10% in Fig. 1B and full list in Supplementary Table 3) and all of these were intrachromosomal. There was a significant enrichment of fusion junctions at chromosomes 17 and 22 (Supplementary Fig. 1A). Furthermore, there was a significant enrichment of fusions that occurred in >10% of tumors at the same chromosomes (Fig. 1B).

**Figure 1.**
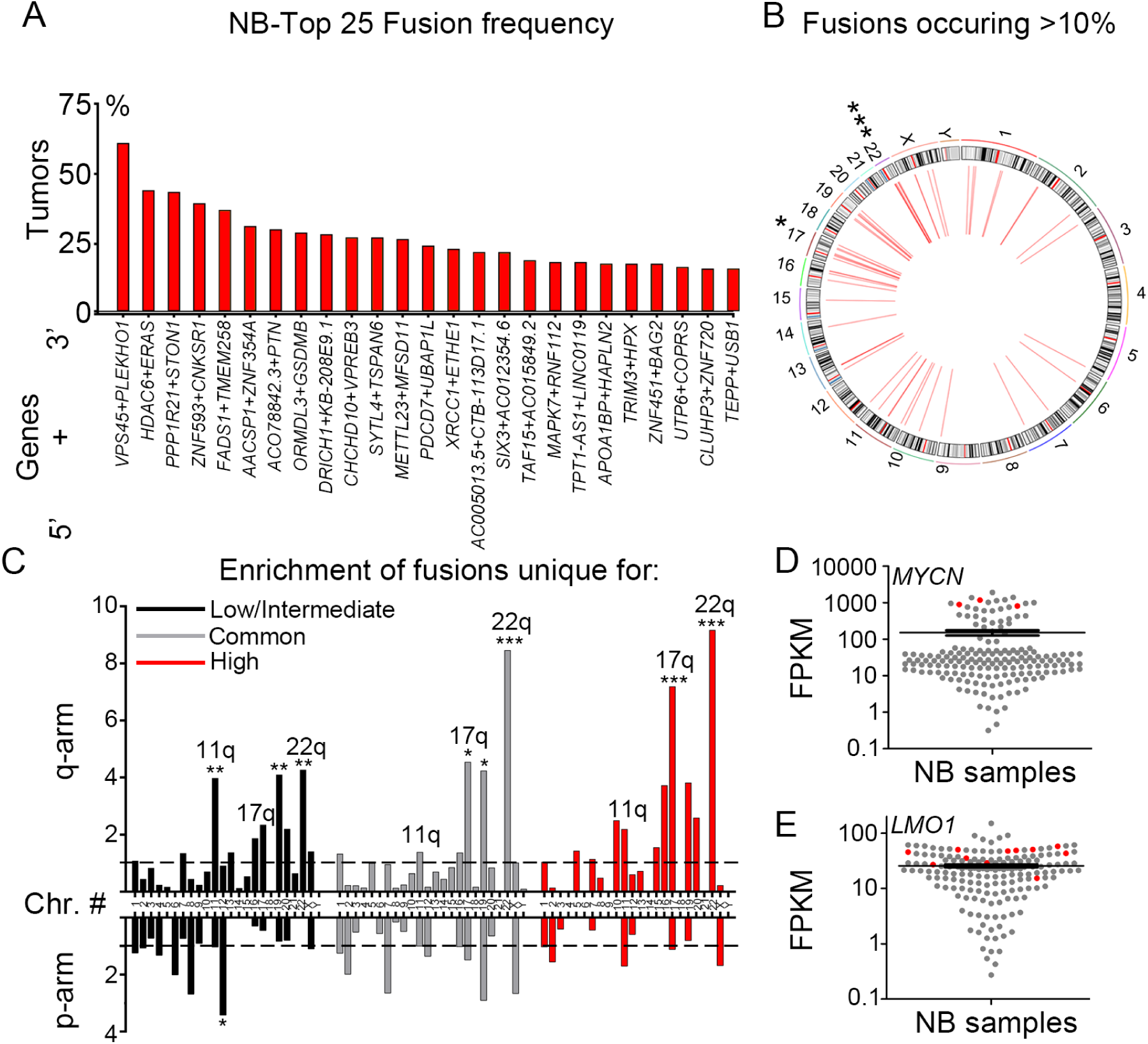
**Identification of fusion transcripts in neuroblastoma tumors.** (A) Top 25 most frequent fusions by FusionCatcher in a cohort of 172 paired end RNA sequenced neuroblastoma patient samples derived from the NCI TARGET project (NB172). (B) Circos plot of genomic distribution of top frequent fusions (> 10%) in NB172. (C) Enrichment of fusion transcripts common or unique to low/intermediate-risk or high-risk tumors in chromosomal arms as calculated by a normalized enrichment score = (counts of fusion transcripts in each chromosomal arm / length of chromosomal arm (Mb)) / average enrichment. (D-E) *MYCN/LMO1* expression levels are high in neuroblastoma tumors (red) bearing fusion transcripts of which one fusion partner is either *MYCN* or *LMO1,* as shown by expression value FPKM (Fragments Per Kilobase per Million mapped reads). P-values in (B-C) were calculated from Z-values, assuming standard normal distribution. *p<0.05, **p<0.01, ***p<0.001.

Gain of 17q is the most frequently occurring genomic alteration in high-risk neuroblastoma and a marker for adverse clinical outcome (*1*), whereas 22q alterations have been reported to be involved in the transition to metastatic and more aggressive neuroblastoma (*20*). We analyzed the fusion transcripts occurring exclusively in low/intermediate-risk and exclusively in high-risk tumors as well as fusion transcripts common to low/intermediate-risk and high-risk levels (Supplementary Table 4-5). Chimeric transcripts unique to low/intermediate-risk tumors exhibited significant enrichment at several chromosomal arms including 11q, a region commonly lost in high-risk neuroblastoma (Fig. 1C). In contrast, both common and high-risk unique fusion transcripts were enriched at 17q and 22q but not at 11q (Fig. 1C), with a pronounced increase in frequency of 17q fusion transcripts in high-risk tumors (Fig. 1C). Thus, with increased risk the frequency of 17q located fusion junctions also increases and was more than seven times higher than the average fusion rate per chromosomal arm. Several fusion transcripts encompassed factors involved in neuroblastoma pathogenesis, including *ARID1B*, *CASZ1*, *HDAC8*, *LMO1, MYCN, BRCA1, TERT* and *PDE6G* (Supplementary Table 6). Notably, tumors harboring fusion transcripts of well-known neuroblastoma oncogenes (e.g. *MYCN* and *LMO1*) also exhibited high expression levels of their wild-type cognates (Fig. 1 D-E). The high-risk susceptibility locus in *LMO1* is significantly associated with MYCN-non amplified high-risk neuroblastoma but not with MYCN-amplified high-risk neuroblastoma (*21*), interestingly the *LMO1-RIC3* fusion (Supplementary Fig. 1B) was exclusively detected in MYCN-non amplified high-risk neuroblastoma (Supplementary Table 6). The *BRCA1-VAT1* fusion was also only detected in MYCN-non amplified high-risk cases; previously it has been shown that copy number amplification of *BRCA1* in NB is restricted to cases lacking MYCN-amplification (*22*) (Supplementary Table 6).

### Validation of fusion transcripts specific for neuroblastoma

To corroborate the fusion transcripts observed in the NB172 dataset in an independent cohort, we performed paired-end RNA-sequencing in an additional cohort containing 14 neuroblastoma patient samples, together with eight neuroblastoma cell lines, NB-validation (NB-v, Supplementary table 7, Materials and Methods). We identified 139 fusions, of which 82 (∼59%) were present in the NB172 dataset (Fig. 2A and Supplementary Table 8). To investigate whether the identified fusions were neuroblastoma specific, we analyzed a cohort of 161 sequenced tissue samples from human normal adrenal glands (*23*). Out of 342 detected fusions in the adrenal gland cohort, only 23 (∼6.7%) were present in the NB172 dataset and only 4 (∼1.2%) of these were present in the NB-v cohort (Fig. 2A and Supplementary Table 9). This enrichment of common fusion transcripts in the neuroblastoma cohorts vs. the adrenal gland dataset was highly significant (chi-square test with Yate’s correction, p-value<0.0001). Fusion transcript associated genes unique to and shared by the two neuroblastoma datasets were enriched at 17q and 2p, two chromosomal regions where gains are closely associated with high-risk neuroblastoma (Fig. 2B). Parametric analysis of gene set enrichment (PAGE) (*24*) comparing high-risk grade 4 tumors with low-risk grade 4s tumors (according to the International Neuroblastoma Staging System, INSS) in the R2 498-SEQC data base (*25*) showed that the NB172/NB-v common genes identified in Fig 2A are enriched in the grade 4 high-risk tumors (Supplementary Fig. 2A). To further investigate whether the identified fusions are distinct for neuroblastoma we analyzed a set of 65 sequenced rhabdoid tumors (National Cancer Institute TARGET, dbGap Study Accession: phs000470.v17.p7) wherein 2055 unique fusion transcripts were detected. However, the overlap with the fusions detected in NB172 dataset was limited to 44 transcripts (∼2.1%) (Fig. 2C). In a cohort of 177 sequenced osteosarcoma tumors (National Cancer Institute TARGET, dbGap Study Accession: phs000468.v17.p7; Fig. 2C) we could detect 1650 unique fusion transcripts but there was no overlap with the NB172 cohort (Fig. 2C). In contrast to fusions detected in neuroblastoma, the majority of detected fusions in rhabdoid tumors and osteosarcoma were inter-chromosomal (Supplementary Fig. 2B). Detected fusions that occur at higher frequencies than 5% are more abundant in neuroblastoma (in 12.3% of tumors) than in rhabdoid tumor (5.7%) and osteosarcoma (0.4%) (Supplementary Fig. 2C). In osteosarcoma there was a considerable number of tumors (∼24.9%, 44/177) harboring fusions predicting substantial deletion of the *TP53* tumor suppressor, disruption of the gene or a truncation at the C-terminal (Fig. 2D, Supplementary Table 10). Notably, *TP53* is one of the most frequently altered genes in osteosarcoma (*26*). As a comparison we could not detect any fusions containing *TP53* in the neuroblastoma tumors (0/172) (Fig. 2D).

**Figure 2.**
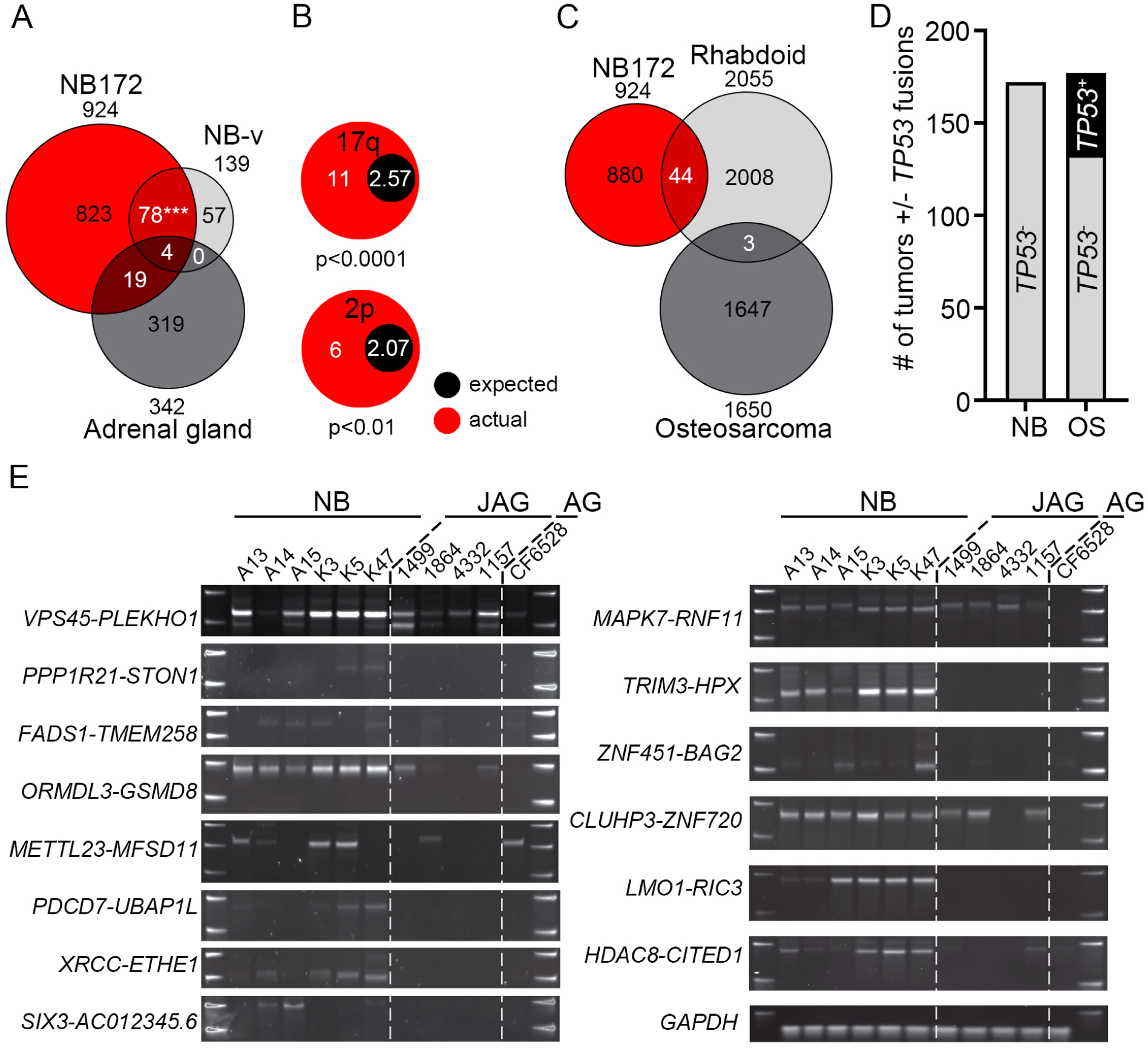
**Validation of fusion transcripts.** (A) Venn diagram of identified fusions in three datasets, NB172 (NCI TARGET), Validation-NB (NB-v, 14 neuroblastoma patients plus 8 neuroblastoma cell lines) and adrenal gland (161 samples from normal human adrenal gland). A significant number of fusions detected in NB-v were present in the NB172 cohort, chi-square with Yate’s correction p-value <0.001. (B) Enrichment of fusion transcript genes unique to the NB172 and the NB-v cohorts on chromosomal arms 2p and 17q. (C) Venn diagram of identified fusions in three datasets, NB172 (NCI TARGET), Rhabdoid (65 Rhabdoid tumor patient samples) and Osteosarcoma (177 Osteosarcoma patient samples). (D) Comparison of the number of neuroblastoma (NB) and osteosarcoma (OS) tumors with at least one fusion transcript containing *TP53*. (E) Validation of fusion transcripts by RT-PCR in neuroblastoma tumors (NB), juvenile adrenal glands (JAG) and adult adrenal gland (AG). *GAPDH* was used for normalization. P-values in (A) were calculated by chi-square test with Yate’s correction and in (B) by Fisher’s exact test. ***p<0.001.

Several programs are available for the identification of fusion transcripts from RNA-seq data. However, results vary considerably between different algorithms (*27*) why validation with independent methods is warranted. We designed primers spanning the fusion junctions of selected chimeric transcripts which occurred at high frequency or encompassed genes previously shown to be important for neuroblastoma biology. We initially designed junction spanning primers for the selected chimeric transcripts *VPS45-PLEKHO1, METTL23-MFSD11, HDAC8-CITED1, ZNF451-BAG2, TRIM3-HPX* and *PRR11-SMG8* (Supplementary Table 11). We performed PCR in 10 neuroblastoma tumor samples and in a panel consisting of cDNA from 14 untransformed human tissues. All selected candidates, except *PRR11-SMG8*, were expressed in a neuroblastoma specific manner (Supplementary Fig. 2D). For validation, PCR-products including three additional fusions, *TAF15-AC015849.2, FADS1-TMEM258* and *LMO1-RIC3*, were excised, inserted into the pCR-Blunt II-TOPO vector and subsequently sequenced. All the sequenced PCR-products exhibited an identical sequence of nucleotides to that of the fusions identified by FusionCatcher (Supplementary Fig. 2D and Supplementary Table 11-12). A previous report identified the *VPS45-PLEKHO1* fusion in non-transformed tissue (*16*), but it was not detected in our panel of non-transformed tissues. Neuroblastoma commonly appear in and around the adrenal gland (*1*). To further investigate specificity of identified fusion transcripts in neuroblastoma compared to healthy juvenile adrenal glands we performed PCR for 14 fusion transcripts on six neuroblastoma tumors, four healthy juvenile adrenal glands (age 0-5 years) and a pool of normal adult adrenal glands (Fig. 2E). The PCR analysis shows that there is an enrichment of fusions in neuroblastoma but also that a subset of normal juvenile adrenal glands expresses certain fusions, albeit at lower levels. Importantly, the PCR validation shows that fusions detected at high frequency with FusionCatcher have a high probability of being true positives.

### The *ZNF451-BAG2* fusion generates a truncated BAG2 protein, present in a subset of neuroblastoma tumors

One of the 25 most frequently occurring fusion transcripts encompassed the *BCL2 associated athanogene* (*BAG2*) which encodes a co-chaperone, BAG2, involved in targeting misfolded proteins for degradation through an ubiquitin independent pathway (*28*). BAG2 levels have previously been shown to increase upon neuronal differentiation in neuroblastoma cells (*29*). In addition, BAG2 clears phosphorylated TAU from neuronal microtubule (*28*), potentially promoting stabilization of axons, which is important for neuronal differentiation. We thus selected *ZNF451-BAG2* for further functional characterization.

The *ZNF451-BAG2* fusion spans the 3’ UTR or exon 14 of *ZNF451* and the second exon of *BAG2,* potentially generating a truncated *BAG2* transcript lacking the first exon (Fig. 3A). Its first exon encodes part of a coiled-coil domain that is absent in the *ZNF451-BAG2* fusion (Fig. 3A). Full length *BAG2* encodes a 23.8 kDa protein, whereas the *ZNF451-BAG2* fusion transcript encodes a smaller 19.6 kDa protein (ΔBAG2) (Fig. 3A). The *ZNF451-BAG2* chimera was present in 31 of the 172 sequenced tumors (18%). Alignment of wild-type BAG2 protein (BAG2) across different species showed that in ΔBAG2 the highly conserved N-terminal coiled-coil domain was truncated (Supplementary Fig. 3A), implying functional relevance of the truncated region for BAG2. To understand if tumors wherein *ZNF451-BAG2* was identified (Supplementary Fig. 2D) also had detectable levels of ΔBAG2 protein, we performed immunoblotting with an antibody targeting BAG2. All probed (n=13) tumors contained BAG2 protein at varying levels however only five tumors also co-expressed detectable levels of ΔBAG2, while no detectable levels of ΔBAG2 were observed in tissue from six human normal adrenal glands (Fig. 3B and Supplementary Fig. 3B).

**Figure 3.**
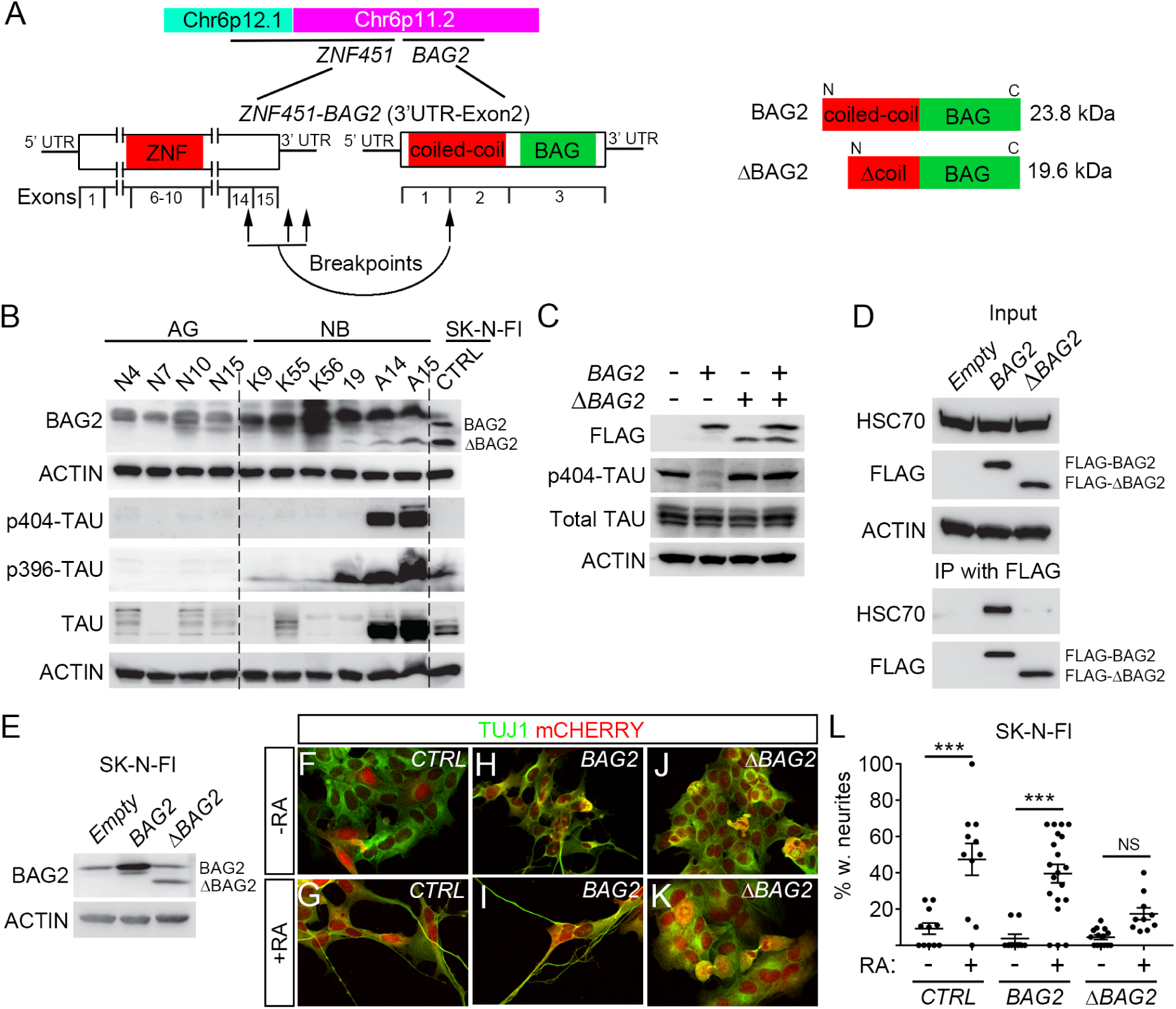
**ΔBAG2 impairs the clearance of phosphorylated TAU and binding to HSC70 and inhibits retinoic acid (RA) induced differentiation in neuroblastoma cells.** (A) Schematic representation of *ZNF451-BAG2* fusion, the resulting truncated BAG2 is referred to as ΔBAG2. (B) Endogenous tumor ΔBAG2 protein expression is associated with high levels of phosphorylated TAU (p-TAU) in neuroblastoma tumors as detected by immunoblotting. (C) SKNFI cells were transfected with p3XFLAG-CMV14-empty/*BAG2*/Δ*BAG2*/*BAG2*+Δ*BAG2* for 48 hours. Cells were harvested and proteins were extracted; whole-cell lysates were used to detect FLAG, p-TAU on Ser404, total TAU and ACTIN. (D) FLAG-tagged proteins were immunoprecipitated from whole-cell lysates as prepared in (C) using Anti-FLAG M2 magnetic beads and eluted. Immunoprecipitated proteins were western blotting to detect HSC70, FLAG2. (E-L) Constitutive lentiviral overexpression of *ΔBAG2*, but not wildtype *BAG2* (backbone pLVX-EF1α-IRES-mCherry) inhibited RA-induced differentiation (6 days of treatment) in SK-N-FI cells. Data in L is represented as mean of transduced cells with TUJ1^+^ neurites/total number of transduced cells +/-SEM, each data-point represents this ratio in a single 10x microscopic field (n=10-20). ***p<0.001, one-way ANOVA with Tukey’s multiple comparison test.

### *ΔBAG2* impairs clearance of phosphorylated TAU and binding to HSC70

BAG2 has been shown to be important for clearance of phosphorylated forms of the TAU protein (pTAU) and thus been implicated as a stabilizer of microtubules (*28*). To test if this capacity was attenuated by the presence of ΔBAG2 we probed the levels of pTAU in a panel of neuroblastoma tumors and normal adrenal glands. This showed a clear association between the presence of pTAU and endogenous ΔBAG2 (Fig. 3B and Supplementary Fig. 3B). To validate that this was caused by ΔBAG2 protein expression, we cloned and validated the *ΔBAG2* transcript whereafter we expressed it and *BAG2* alone or in combination in SK-N-FI neuroblastoma cells. Upon *BAG2* overexpression, the levels of pTAU were significantly reduced whereas total TAU was present in amounts similar to those in control-transduced cells (Fig. 3C). *ΔBAG2* overexpressing cells retained pTAU levels and more importantly, upon co-expression BAG2 failed to clear pTAU (Fig. 3C), implying that ΔBAG2 can act as a negative regulator of BAG2 function. BAG2 has been shown to bind the heat shock cognate 70 (HSC70) (*30*), a chaperone protein important for pTAU ubiquitination (*31*) and axon outgrowth (*32*). Overexpression of *BAG2* and *ΔBAG2* in SK-N-FI neuroblastoma cells followed by BAG2 immunoprecipitation via FLAG revealed that BAG2 but not ΔBAG2 binds a 70 kDa protein (Supplementary Fig. 3C). Since BAG2 has been reported to bind HSC70 (*30*), we performed additional immunoprecipitation followed by immunoblotting with a HSC70 specific antibody. This revealed that BAG2 binds HSC70, whereas ΔBAG2 does not (Fig. 3D).

### *ΔBAG2* impedes differentiation of neuroblastoma in response to retinoic acid

To investigate whether ΔBAG2 impinges on the capacity of neuroblastoma cells to differentiate, we treated *CTRL* (empty vector)*, BAG2* or *ΔBAG2* expressing SK-N-FI cells (Fig. 3E) with retinoic acid (RA), a compound used in adjuvant therapy of high-risk neuroblastoma patients (*17*). After six days of RA treatment, cells transduced with either *CTRL* (Fig. 3F-G, L) or *BAG2* (Fig. 3H-I, L) acquired neuronal morphology with long neurites. In contrast, *ΔBAG2* transduced cells exhibited a weaker response to RA, with significantly less neurite formation (Fig. 3J-L). To validate this effect we transduced SK-N-BE(1) neuroblastoma cells with a doxycycline-inducible version of the *ΔBAG2* fusion genes (Supplementary Fig. 3D). Upon doxycycline induction, *ΔBAG2* expressing cells exhibited a reduced capacity to respond to RA (Supplementary Fig. 3E-I).

### Fusion transcripts with splicing motifs are enriched in neuroblastoma and elevated expression of spliceosome factors is a strong predictor of high-risk tumors

Previously it has been suggested that *cis*-splicing between adjacent genes can generate fusion transcripts in prostate cancer (*7, 10*). To elucidate whether there was a correlation between distance and frequency we plotted the distance between 5’ and 3’ of the fusion junction in the intrachromosomal fusion transcript versus the fusion frequency in the NB172 and NB-v data sets (Fig. 4A). Fusions occurring at high frequency in both data sets were enriched between 1 to 100kb, whereas transcripts separated by more than 100kb occurred at lower frequencies and almost exclusively in the larger NB172 data set (Fig. 4A). The genomic proximity of identified transcripts suggests that these fusions could be a result of *cis*-splicing (*7, 12*). Inspection of nucleotide sequences located 5’ and 3’ of the fusion junctions for canonical splicing donors (GT) and acceptors (AG) revealed that 81.9% carried GT and AG at the 5’ and 3’ fusion sites (GT*AG) (Fig. 4B). Notably, 87% of intra-chromosomal fusions had GT*AG at the fusion junction whereas only 45.9% of inter-chromosomal fusions contained canonical splicing sites at the junctions. Thus, in total there was ∼3 times more intrachromosomal fusions with canonical splicing sites than expected by chance in neuroblastoma, (p-value<0.0001, chi-square test with Yate’s correction). This pattern was unique to neuroblastoma as fusions detected in normal adrenal gland, osteosarcoma and rhabdoid tumor exhibited no enrichment of intrachromosomal fusions with splicing sites (Fig. 4B). Aberrant RNA splicing has been suggested as a driving event for several cancers and mutations in genes coding for components of the spliceosome have been identified in several tumors (*33*). Notably, in *MYC*-driven breast cancer, it has been shown that spliceosome is a therapeutic vulnerability (*4*), whereas in small cell lung cancer spliceosome inhibition was reported to be effective regardless of *MYC* status (*34*). We compared expression levels of genes of the KEGG, “Spliceosome” gene category between neuroblastoma and normal adrenal gland. Out of 134 genes 46 had significantly higher expression levels in the neuroblastoma, whereas only three had lower levels of expression (Fig. 4C). To investigate whether there were clinically relevant differences in expression of spliceosome genes between low- and high-risk neuroblastoma we performed k-means analysis of the 498-SEQC neuroblastoma cohort (Fig. 4D). Together with previous observations (*35*) our analysis elucidates how the differential expression of spliceosome factors clearly identifies tumors of different clinical outcome, with high expression levels of splicing factors predicting high-risk tumors with bleak clinical outcome and substantially shorter overall survival (Fig. 4D-E). Furthermore, inspection of previous genome sequencing studies of neuroblastoma patients revealed mutations in several spliceosome factors in primary tumors and *de novo* mutations in spliceosome factors occurred in relapsed tumors (*2, 3, 36, 37*) (summarized in Supplementary Table 13).

**Figure 4.**
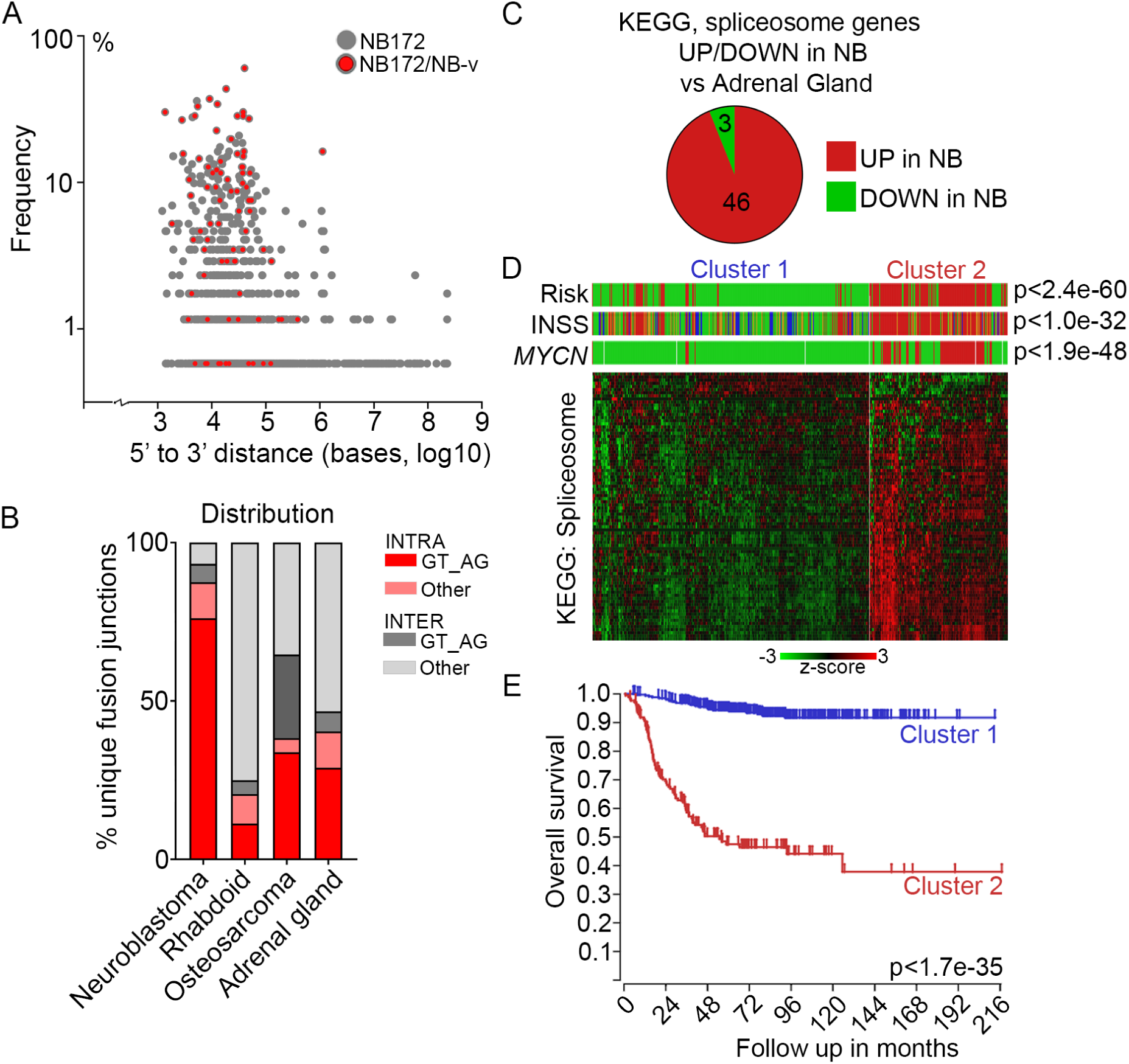
**Enrichment of canonical splicing pattern at fusion junctions is associated with aberrant spliceosome activity in high-risk neuroblastoma.** (A) Plot of the frequency versus the distance from 5’ to 3’ in the intrachromosomal fusion transcript identified in the NB discovery cohort (NB172); each dot represents a unique fusion transcript; fusion transcripts re-identified in the NB validation cohort (NB-v) were marked as red. (B) Distribution of unique intrachromosomal vs interchromosomal fusion junctions flanked by GT_AG or other nucleotide motifs in neuroblastoma, rhabdoid tumor, osteosarcoma and normal adrenal gland. (C) Differential expression of genes in the KEGG Spliceosome pathway between neuroblastoma dataset (NBL172) versus human normal adrenal gland dataset. (D) K-means analysis of the 498-SEQC dataset, utilizing the genes in the KEGG spliceosome pathway, generates two clusters with significant differences in Risk, INSS and *MYCN*-amplification. (E) Overall survival probability of the two clusters identified in (D).

### Spliceosome inhibition impedes expression of fusion transcripts, induces neuroblastoma apoptosis and inhibits growth of neuroblastoma xenografts

To investigate whether inhibition of spliceosome activity would selectively impede the generation of fusion transcripts but not their wild type cognates, we treated neuroblastoma cells (LAN-1 and SK-N-BE(1)) with the spliceosome inhibitor, Pladienolide B, targeting SF3B1 and PHF5 key components of the splicing machinery (*38, 39*). Upon treatment, there was a loss of expression for a majority of selected high frequency fusion transcripts whereas expression of wild type genes constituting the fusions were largely unaffected at these concentrations (Fig. 5A-B and Supplementary Fig. S4). To elucidate if selective loss of fusion transcripts upon spliceosome inhibition was associated with increased apoptosis we treated the neuroblastoma cell lines LAN-1, SK-N-BE(1) and SK-N-AS with increasing concentrations of Pladienolide B. As controls we utilized a panel of transformed and non-transformed human cells consisting of U87 (glioma), U2OS (osteosarcoma) and NHA (normal human fibroblasts). Already at 5nM the neuroblastoma cell lines exhibited increased levels of cleaved caspase-3 and cleaved PARP, whereas control cells showed no signs of cell death. At 10nM cell death in both neuroblastoma cell lines was further increased but control cells were still non-responsive (Figure 6A). We repeated these experiments with SRPIN340 (*40*), an inhibitor of SRPK1 which phosphorylates proteins containing serine-arginine-rich (SR) domains, a function necessary for proper splicing. As with Pladienolide B, SRPIN340 selectively induced apoptosis in neuroblastoma cells but not in the other cell types tested (Fig. 6B). To test whether inhibition of the spliceosome could affect neuroblastoma growth *in vivo* we transplanted LAN-1 cells into the flanks of nude mice. When the xenografts had reached a volume of approximately 200mm^3^ we initiated treatment with Pladienolide B. At the endpoint, Pladienolide B treated tumors exhibited significantly reduced growth in comparison to vehicle treated tumors (Fig. 6C-D).

**Figure 5.**
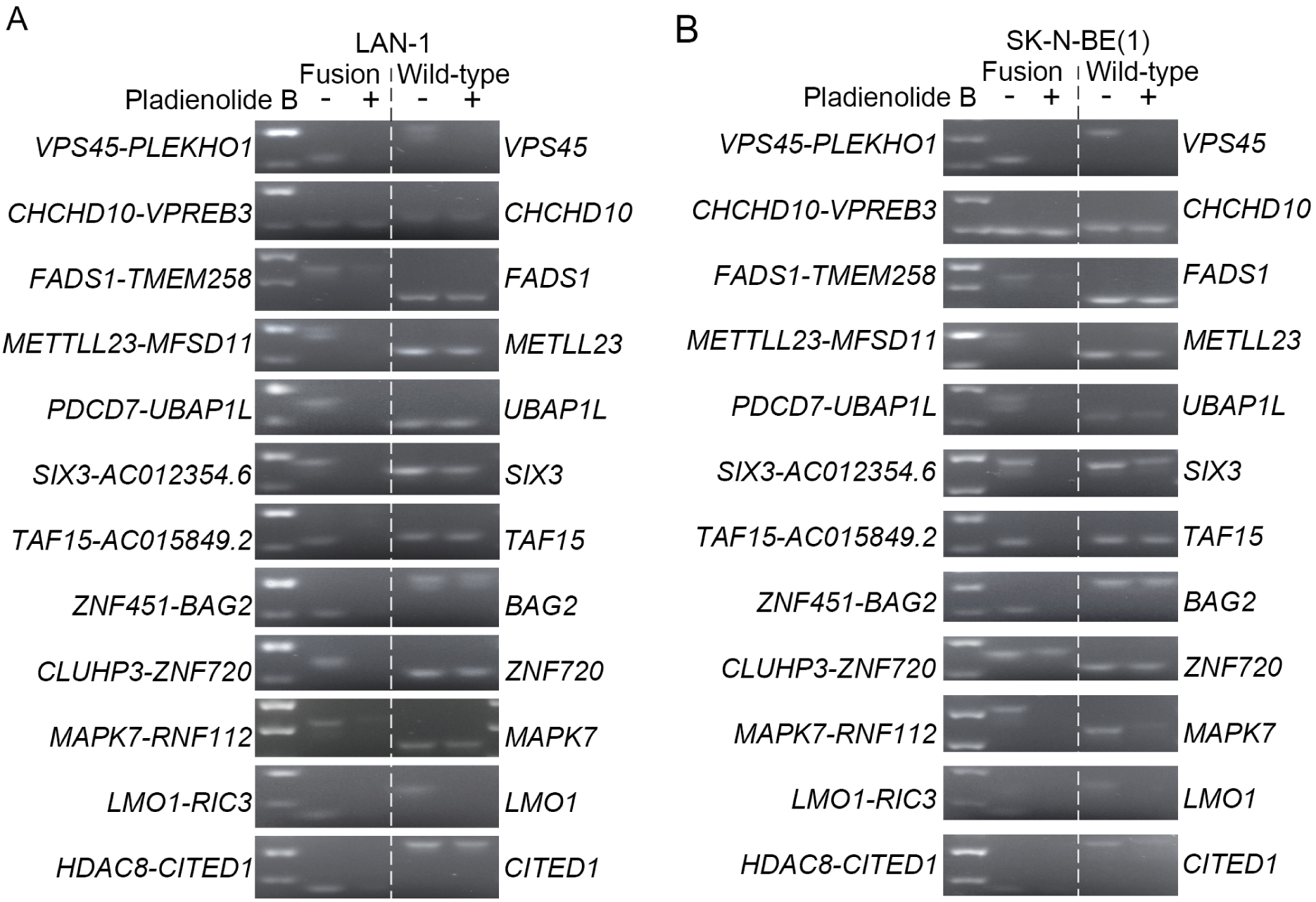
**Pharmaceutical inhibition of spliceosome activity selectively reduces expression of fusion transcripts in neuroblastoma cells.** (A-B) Splicing-dependent generation of several high frequent fusion transcripts in LAN-1 (A) and SK-N-BE(1) (B) neuroblastoma cells. SK-N-BE(1) and LAN1 cells were treated with splicing inhibitor Pladienolide B at 100 nM (SK-N-BE(1)) and 5-50 nM (LAN-1) for 6 hours; RNA was isolated from harvested cells and reversely transcribed to cDNA; RT-PCR were performed with primers spanning the fusion junctions (fusion transcript) or exon-exon boundary (wild-type cognate).

**Figure 6.**
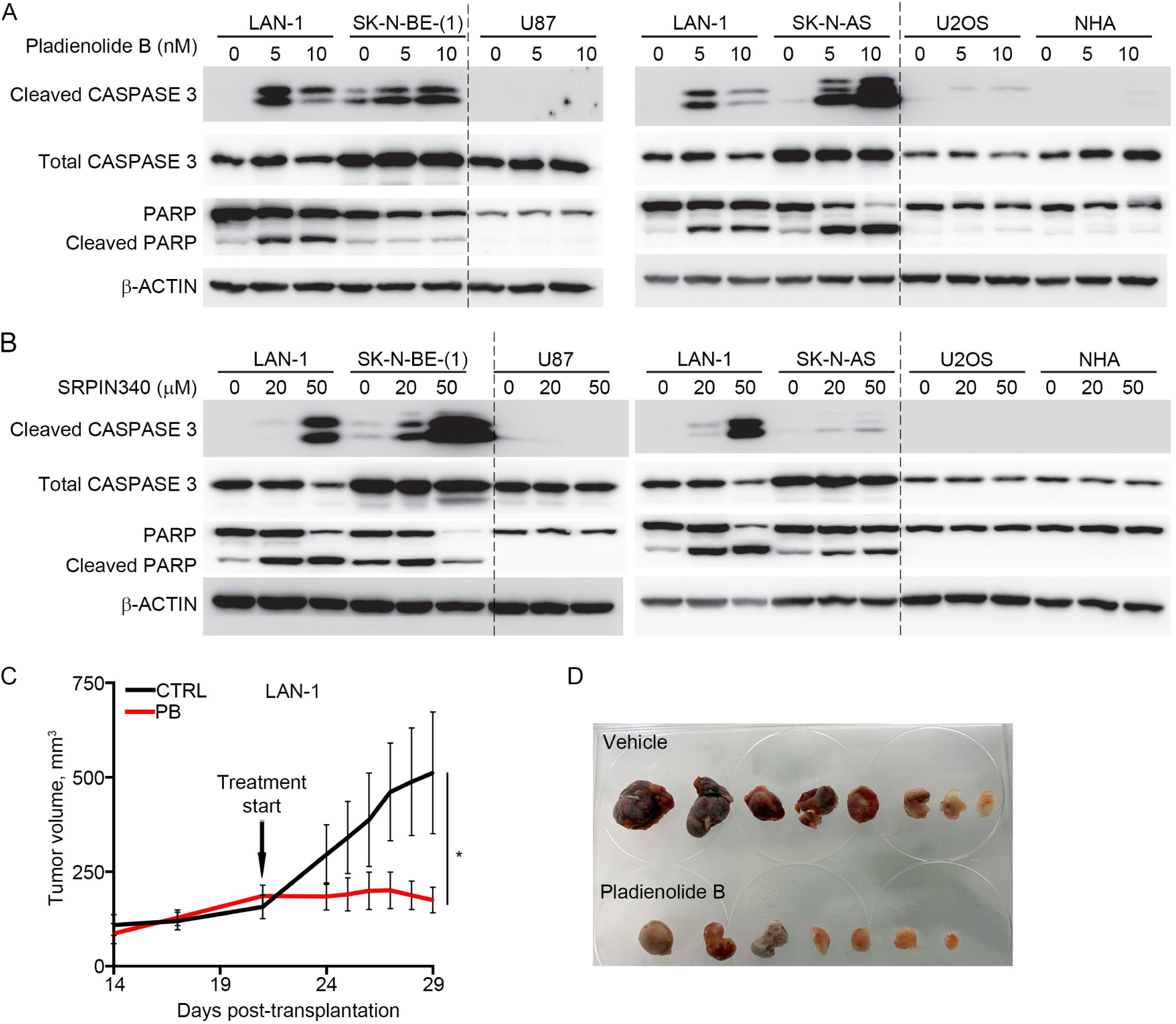
**Pharmaceutical inhibition of spliceosome activity selectively induces apoptosis in neuroblastoma cells and impedes tumor growth in neuroblastoma xenografts.** (A) Induction of apoptosis in LAN-1,SK-N-BE(1) and SK-N-AS neuroblastoma cells, but not in human glioma cells (U87), osteosarcoma cells (U2OS) or normal human astrocytes (NHA) upon pladienolide B treatment (0-20 nM) for 48 hours, as detected by western blotting of full-length CASPASE-3, cleaved CASPASE-3, full-length PARP and cleaved PARP; β-ACTIN was used as a loading control. (B) Induction of apoptosis in LAN-1,SK-N-BE(1) and SK-N-AS neuroblastoma cells, but not in human glioma cells (U87), osteosarcoma cells (U2OS) or normal human astrocytes (NHA) upon SRPIN340 treatment (0-50 mM) for 48 hours, as detected by western blotting of full-length CASPASE-3, cleaved CASPASE-3, full-length PARP and cleaved PARP; β-ACTIN was used as a loading control. LAN-1 cells were included in all western blots (A-B) for comparison. (C-D) In mice xenografted with LAN-1 neuroblastoma cells, treatment with Pladienolide B significantly reduced tumor growth (n=7-8) (C-D). *p<0.05; Mixed-effects model with Geisser-Greenhouse’s epsilon correction.

## Discussion

Our analysis revealed that high expression levels of splicing associated factors is a distinguishing feature of high-risk neuroblastoma, representing a strong predictor of poor clinical outcome. Furthermore, pharmacological inhibition of the spliceosome selectively induced apoptosis in neuroblastoma cells and abrogated tumor growth *in vivo*. Our study shows that a high frequency of neuroblastoma specific fusion transcripts could constitute an overlooked process through which altered transcripts are generated. It has been shown that upon cellular stress (e.g. viral infection, replicative or osmotic stress and mutational events) transcriptional termination can be blocked, increasing the probability of generating “downstream of genes”-transcripts (*15*). Thus, a proportion of fusions could represent passengers that occur as a response to cellular stress combined with neuroblastoma associated events such as gain or loss of chromosomal regions (e.g. 17q and 2p). In contrast to such passengers, certain fusions could be early events preceding and promoting other transformative events. Pharmacologic inhibition of splicing selectively repressed expression of several top frequent fusion transcripts but not of their wild-type cognates and there was a high frequency of splicing donor/acceptor sites in neuroblastoma specific fusions but not in fusions detected in normal adrenal gland, osteosarcoma or rhabdoid tumors. This pattern implies that a substantial proportion of the detected fusions are of the same *cis*-splicing type as reported in prostate cancer (*7*). Regardless of the mechanisms underlying the generation of these fusions, they are not dependent on amino acid changing mutations but can still provide a source of modified gene products with a potential to promote neuroblastoma but also reveal novel drug targets. Given the low frequency of recurrent mutations in neuroblastoma, such a pool of altered gene products could indeed be relevant for tumor pathophysiology. A background of expressed fusion transcripts could potentially augment the effect of oncogenic drivers. Notably, our analysis shows that when established drivers of neuroblastoma (e.g. *MYCN* and *LMO1*) are part of the fusion transcripts the expression levels of wild type cognates are elevated. In addition, a panel of neuroblastoma specific fusions occurring at high frequency could serve as biomarkers for diagnosis and the presence of risk-specific fusions could sub-divide neuroblastoma patients for precision therapy. A previous study indeed shows how the *SLC45A3-ELK4* read-through fusion transcript is elevated in prostate cancer tissue, is androgen-regulated and can be detected in an non-invasive assay from biopsies of men at risk of having prostate cancer (*13*).

One concern with previously reported fusions is that relatively few cases have been independently validated. In 2015, only 3% of fusions identified by deep sequencing could be reproducibly detected (*9*). Arguably, the “non-genomic” characteristic of this type of intrachromosomal fusions potentially augments the detection of false positives. The risk that a proportion of fusion transcripts constitute false positives as the result of spurious transcription or of sequencing errors is reduced by our crosswise analysis with other tumors and healthy tissues. There is a clear enrichment of common fusions unique for the neuroblastoma data sets that do not appear in any of the other tumors nor in adult normal adrenal glands. Neuroblastoma specific fusions are enriched for genes located at chromosomal regions (2p or 17q) which are commonly gained in high-risk neuroblastoma. PCR with primers spanning the fusion points and selective loss of several fusion transcripts upon spliceosome inhibition provide support, independent of bioinformatic tools, that splicing dependent fusion transcripts indeed are present. Interestingly, in a few of the examined juvenile adrenal glands there was low but detectable expression of certain fusion transcripts. Thus, our data suggest that certain fusion transcripts can be lowly expressed at the most common site of neuroblastoma and then be enriched for upon tumor formation. Tumor specific distribution of fusions is also evident in the osteosarcoma tumors where the *TP53* tumor suppressor is a fusion partner in an disproportional amount of the detected fusions, reflecting the importance of inactivating mutations in *TP53* for this disease (*26*). It has previously been reported that the presence of short homology sequences (SHS) can generate false RNA-chimeras due to template switching during the reverse transcriptase reaction (*19*). Such a mechanism would presumably generate random fusions between transcripts containing matching SHSs with no preference for any particular chromosomal location nor for intrachromosomal vs interchromosomal fusion transcripts. The non-random enrichment of fusions at chromosomal locations that mirror the disease as well as the enrichment of intrachromosomal fusions between closely located genes suggests that reverse transcriptase induced template switching is not the cause of these fusions. Moreover, our study shows that an altered protein with novel properties can be generated as a consequence of an intrachromosomal, splicing dependent fusion. This altered protein influences the response to a drug (RA) commonly used to treat high-risk neuroblastoma patients.

Even though our study provides novel insights of the role of splicing and splicing-dependent RNA-fusions in neuroblastoma, future studies are necessary to functionally test whether the selective loss of RNA-fusion expression and impeded tumor growth upon spliceosome inhibition are causally linked. In addition, the functional role of distinct fusions as well as the total impact of collected fusions remain to be investigated.

Taken together, our data show that high expression levels of splicing associated factors are characteristic of high-risk neuroblastoma and that the spliceosome can serve as a target for pharmacological intervention. In addition, splicing dependent RNA-fusions provide a mechanism, distinct from amino acid changing mutations, through which altered gene products can be generated. This could potentially compensate for the lack of recurrent mutations in NB. Enrichment of fusions at chromosomal regions associated with NB and the specificity of expression imply that this type of altered gene products are relevant for the biology of the disease and could serve as potential novel drug targets and diagnostic markers.

## Materials and Methods

### Ethical considerations

All animal experiments were performed according to Swedish guidelines and regulations, the ethical permit 6420-2018 was granted by “Stockholms Norra djurförsöksetiska nämnd, Sweden”. Neuroblastoma primary tumors came either from the Swedish NB Registry (ethical permission (DN03-736) granted by Karolinska Institutets Forskningsetikommitté Nord, (clinical information described in Li et. al. (*41*), or from the Irish NB cohort (described in Supplementary table 7), with ethical approval of the Medical and Research Ethics Committee of Our Lady’s Children’s Hospital, Crumlin, Dublin, Ireland. Informed consent from families of subjects was obtained for samples. Six histologically confirmed normal human adrenal glands were included as controls (covered by existing ethical approvals; Dnr 01-136 and Dnr 01-353 KI forskningsetikkommitté Nord). Four juvenile adrenal glands were from University of Maryland Brain and Tissue Bank; Total RNA of pooled normal adrenal gland was from Clontech (Cat# 636528). Human total RNA from different normal tissues from Clontech (Human Total RNA Master Panel II,Cat#. 636643).

### RT-PCR and Paired-end RNA-seq

RNA was isolated using the PerfectPure RNA Cultured Cell Kit (cell lines) and PerfectPure RNA Tissue Kit (patient samples) from 5 PRIME. Reverse transcription was performed with iScript cDNA Synthesis Kit (Bio-Rad) and RT-PCR was performed by using iQ SYBR Green Supermix (Bio-Rad). RNA-seq libraries were prepared using TruSeq RNA Library Preparation Kit v2 (Illumina); paired-end RNA sequencing (125 bp) were performed in SciLifeLab (Stockholm).

### Data Analysis

FusionCatcher (*18*) was applied to detect the fusion transcripts in paired-end RNA-seq data. Reads were mapped to hg19 and differential expression analysis was performed as described in (*42*). For enrichment analysis in Figure 1C, only fusions occurring above ∼3% in each risk group were included (1 case in Low/Intermediate-risk, 4 cases in High-risk and 5 cases for the common fusion transcripts shared by Low/Intermediate-risk and high-risk groups). For the Z-test in 1C the μ and σ values were calculated from genome wide enrichment of fusions at the different chromosomal arms.

### Cloning and expression of wildtype *BAG2* and *ZNF451-BAG2*

cDNA was synthesized from total RNA by the iScript cDNA Synthesis Kit (Bio-Rad). Coding regions of wildtype *BAG2* and *ZNF451-BAG2* were amplified from SK-N-AS cells for subcloning into p3XFLAG-CMV14 (Sigma) and pLVX-EF1α-IRES-mCherry (Clontech) using primer pair 1 and 2 respectively (Supplementary Table 11), inserted to pCR-Blunt II-TOPO vector via zero Blunt TOPO PCR Cloning Kit (ThermoFisher Scientific) and sequenced. *BAG2* and *ZNF451-BAG2* were subcloned into p3XFLAG-CMV14 vector using EcoRI and BamHI and pLVX-EF1α-IRES-mCherry lentiviral vector using *Eco*RI and *Bam*HI. To generate pLVX-Tetone-puro-IRES-mCherry-empty/BAG2/ZNF451-BAG2 vectors, pLVX-Tetone-puro-empty (Clontech) construct was linearized with *Age*I, blunted and digested with *Eco*RI; pLVX-EF1α-IRES-mCherry-empty/BAG2/ZNF451-BAG2 vectors were linearized with *Mlu*I, blunted and digested with *Eco*RI to release IRES-mCherry-empty/BAG2/ZNF451-BAG2; then linearized and blunted pLVX-Tetone-puro-empty construct was ligated with fragment of IRES-mCherry-empty/BAG2/ZNF451-BAG2 separately. Lentiviruses expressing pLVX-EF1α-IRES-mCherry-empty/BAG2/ZNF451-BAG2 and pLVX-Tetone-puro-IRES-mCherry-empty/BAG2/ZNF451-BAG2 were produced in 293FT cells using lipofectamine 2000 based protocol.

### Cell culture

All neuroblastoma cell lines (SH-SY5Y, SK-N-SH, SK-N-FI, SK-N-BE(1), SK-N-AS, LAN-1, CHP-212 and IMR-5) were maintained in RPMI1640 medium supplemented with 10% FBS, 1% penicillin/streptomycin and 1% L-glutamine and grown in 5%CO2 at 37°C.

### Differentiation assay, immunofluorescence staining and apoptosis analysis

For short-term RA induced differentiation assay, SK-N-FI cells were seeded in 6 well plates; after 24 hours, cells were infected by lentiviruses expressing pLVX-EF1-IRES-mCherry-empty/BAG2/ZNF451-BAG2 for 48 hours; then cells were trypsinized and 20,000 cells were reseeded into 6 well plates with coverslips; 24h later, cells were treated with 1 M retinoic acid (RA) or DMSO as control for 6 days. For doxycycline-inducible system, 20,000 SK-N-BE(1) cells with stable overexpression of pLVX-Tetone-puro-IRES-mCherry-ZNF451-BAG2 were seeded in 6 well plates with coverslips, and after 24h, cells were pre-treated with 1 M doxycycline or DMSO as a control for one day; then cells received one of the following 4 different treatment for 4 days: DMSO, 1 M RA, 1 M doxycycline or 1 M RA+1 M doxycycline.

Cells on coverslips were fixed in RPMI1640 medium containing 2% PFA for 5 minutes at room temperature (RT), washed once with PBS at RT, fixed with 4% PFA for another 15 minutes at RT and washed twice with cold PBS; then cells were permeabilized with PBS containing 0.25% Triton X-100 for 15 minutes at RT and blocked with 3% BSA for 1h at RT. Immunofluorescent staining were performed with the following primary antibody Tuj1 (Covance, 1:1000) overnight at +4 C.

For apoptosis analysis, cells were seeded on 6 cm dishes 24 hours before treatment with Pladienolide B (Santa Cruz) or SPRIN340 (MedChemExpress); DMSO was used as a control; after 48 hours treatment, cells, including detached cells in the supernatant, were harvested and total proteins were isolated by lysing cells in lysis buffer containing 100 mM Tris-HCl, pH 7.4, 150 mM NaCl, 1% NP40. Protease and phosphatase cocktail inhibitors (Roche) were added to the lysis buffer before use. Protein levels of apoptosis markers were analyzed by western blotting.

### Xenografts

2 × 10^6^ LAN-1 cells in 200 μL PBS:matrigel (BD, 354248) (1:1) were injected s.c. into the right flank of adult female Crl:Nu (Ncr)-Foxn1 nu (nude mice). Treatments started 14 days after transplantation. The control group was injected i.p. with 100 µL vehicle (10% EtOH and 4% tween80 in sterile PBS) per mouse per day. Pladienolide B was injected i.p. at a concentration of 10mg/kg. Tumors were measured externally using a digital caliper; the greatest longitudinal diameter (length) and the greatest transverse diameter (width) were determined, and tumor volume was calculated using the modified ellipsoidal formula [(length × width^2^)/2]. Overall health was monitored during treatment, and weight was measured weekly. The size of the treated groups was determined based on previous xenograft experiments (*42*). For tumor growth comparison in Fig. 6C, mixed-effects analysis with Geisser-Greenhouse’s epsilon correction was performed. All statistical analysis was performed in Excel or GraphPad Prism 8.1.1.

### Quantification

Cells were stained as in Fig. 4G-L and O-R. Images were taken by confocal microscopy (10X). For quantification, images were coded and a researcher who had not participated in staining and image acquisition manually counted the ratio of transduced cells extending TUJ1^+^ neurites/total number of transduced cells/microscopic field. For DOX^-^ SK-N-BE(1) the ratio of cells extending TUJ1^+^ neurites/total number of cells/microscopic field were counted. Microscopic fields containing less than three transduced cells were discarded. Grubb’s test was performed for outlier detection (alpha p<0.05). For statistical analysis in Fig. 4L and R one-way ANOVA followed by a Tukey’s multiple comparison test was performed.

### Western blotting

Immunoblotting was performed using standard protocols. Following primary antibodies were used: BAG2(sc-390262, 1:200), phospho-ser396-Tau(sc-101815, 1:1000), HSC70(sc-7298, 1:200) from Santa Cruz; phosphor-ser404-Tau(44-758G, 1:200) from ThermoFisher Scientific; total Tau(A0024, 1:1000) from Dako; beta-actin(AC-15, 1:3000), FLAG(F1804, 1:1000) from Sigma; PARP (9542s), Caspase-3 (14220s) and cleaved caspase-3 (9664s) from Cell Signaling.

### Co-immunoprecipitation

SK-N-FI cells were seeded in 15 cm dishes 24 hours before transfection. Transfection of p3XFLAG-CMV14-empty/BAG2/ZNF451-BAG2 constructs was performed using lipofectamine 2000. 48 hours post-transfection, cells were harvested and proteins were extracted using lysis buffer containing 10 mM TRIS-HCl (pH7.4), 150 mM NaCl, 0.5% NP40, complete protease and phosphatase inhibitors (Roche). FLAG-tagged fusion proteins were immunoprecipitated using Anti-FLAG M2 magnetic beads (Sigma) according to manufacturer’s instructions. Bound proteins were examined by silver staining or Western blotting using standard protocols.

## Acknowledgements

We thank members of the Holmberg and Schlisio groups for valuable comments.

## Author contributions

Y.S. and J.H. designed the study. Y.S. performed the majority of the experiments with help from V.R., E.M., S.L., P.B., J.Y., and I.W. Y.S. generated the libraries for RNA-sequencing. Y.S. and J.H performed the analysis with help from O.B.R. C.C.J, A.S., C.L., P.K. and M.J.S supplied the clinical material. Y.S. and J.H. wrote the manuscript with input from all authors.

## Data and materials availability

Data needed to evaluate the conclusions in the paper are present in the paper and/or the Supplementary Materials. Raw and processed NGS data will be deposited in the GEO database.

## Supplementary figure and table legends

**Figure S1.**
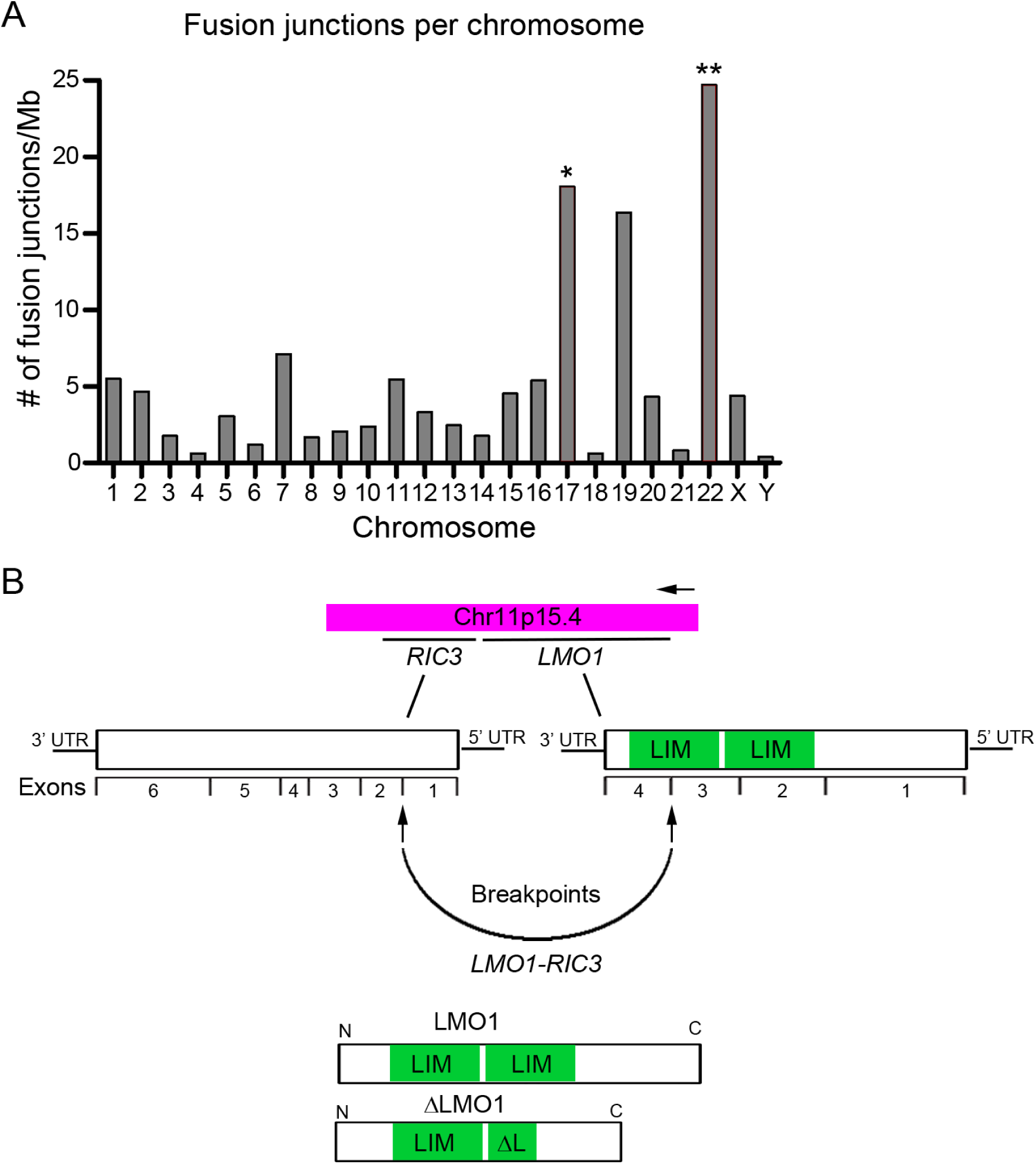
**Identification of fusion transcripts in neuroblastoma tumors.** (A) Genomic distribution of fusion junctions identified by FusionCatcher in a cohort of 172 paired end RNA sequenced neuroblastoma patient samples derived from the NCI TARGET project (NB172). (B) Schematic representation of *the RIC3-LMO1* fusion transcript. P-values in (A) were calculated from Z-values, assuming standard normal distribution. *p<0.05, **p<0.01.

**Figure S2.**
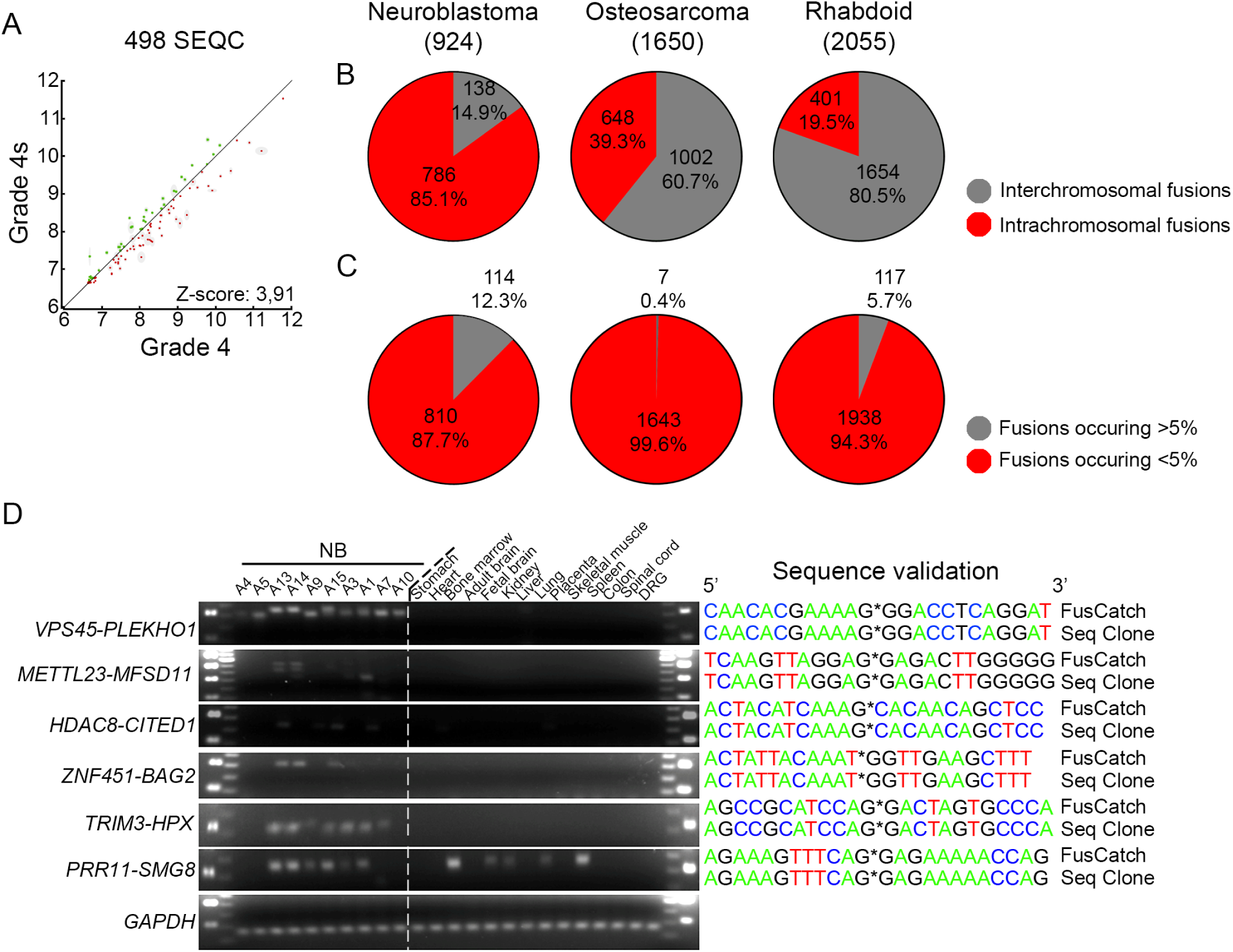
**Comparison of fusions in neuroblastoma, osteosarcoma and rhabdoid tumor and validation of fusions with PCR.** (A) Parametric analysis of gene set enrichment (PAGE) of the 498-SEQC neuroblastoma dataset show an enrichment of NB172 and NB-v unique common fusions in high-risk grade 4 tumors compared to low-risk 4S tumors. (B) Ratio of intra- vs interchromosomal fusion detected in neuroblastoma, osteosarcoma and rhabdoid tumors. (C) Ratio of fusions occurring in more or less than 5% of the patients for neuroblastoma, osteosarcoma and rhabdoid tumors. (D) Validation of fusion transcripts by RT-PCR and sequencing in Validation-NB neuroblastoma patients, indicated normal tissues were used as controls; DRG, human dorsal root ganglion.

**Figure S3.**
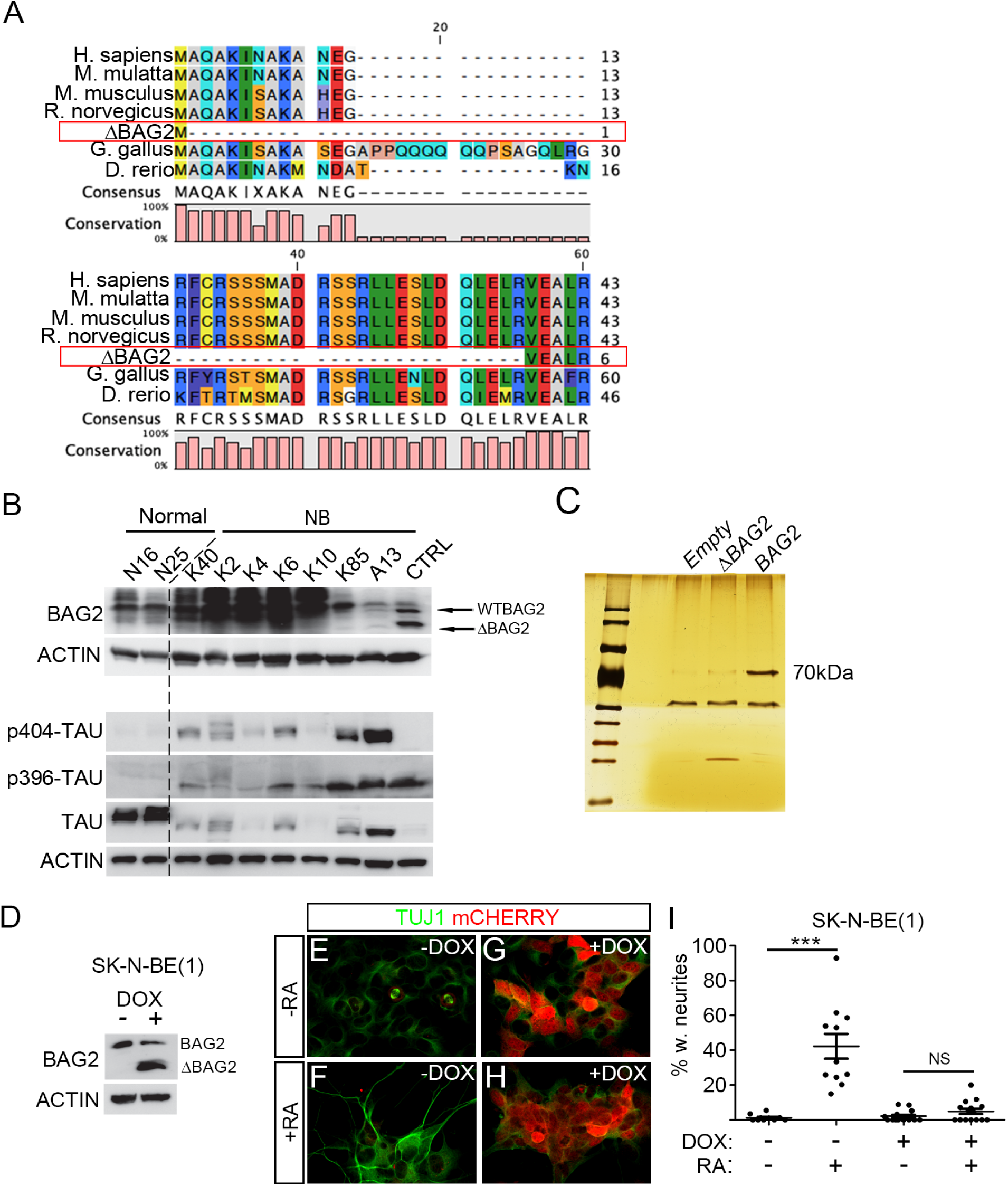
**ΔBAG2 impairs the clearance of phosphorylated TAU and binding to HSC70 and impedes retinoic acid-induced differentiation.** (A) Alignment of ΔBAG2 protein with wildtype BAG2 in different species. (B) Detection of ΔBAG2 protein in additional neuroblastoma tumor samples. N16 and N25 are normal human adrenal glands; other tissues are neuroblastoma patient samples; positive control is the same as in Figure 3B. ΔBAG2 is associated with high levels of phosphorylated TAU in neuroblastoma patients as detected by immunoblotting. (C) FLAG-tagged proteins were immunoprecipitated from whole-cell lysates as prepared in (Fig. 3C) using Anti-FLAG M2 magnetic beads, eluted and then analyzed by silver staining. (D-I) Doxycycline inducible lentiviral overexpression of *ΔBAG2* (backbone pLVX-Tet-one-puro-IRES-mCherry) inhibited RA-induced differentiation (4 days of treatment) in SK-N-BE(1) cells. Protein levels of BAG2 and ΔBAG2 were analyzed by Western blotting (D). Immunostaining was performed using antibody against neuronal marker β3-tubulin (TUJ1). Data in I is represented as mean of transduced cells with TUJ1^+^ neurites/total number of transduced cells +/-SEM, each data-point represents this ratio in a single 10x microscopic field (n=10-20). ***p<0.001, one-way ANOVA with Tukey’s multiple comparison test.

**Figure S4.**
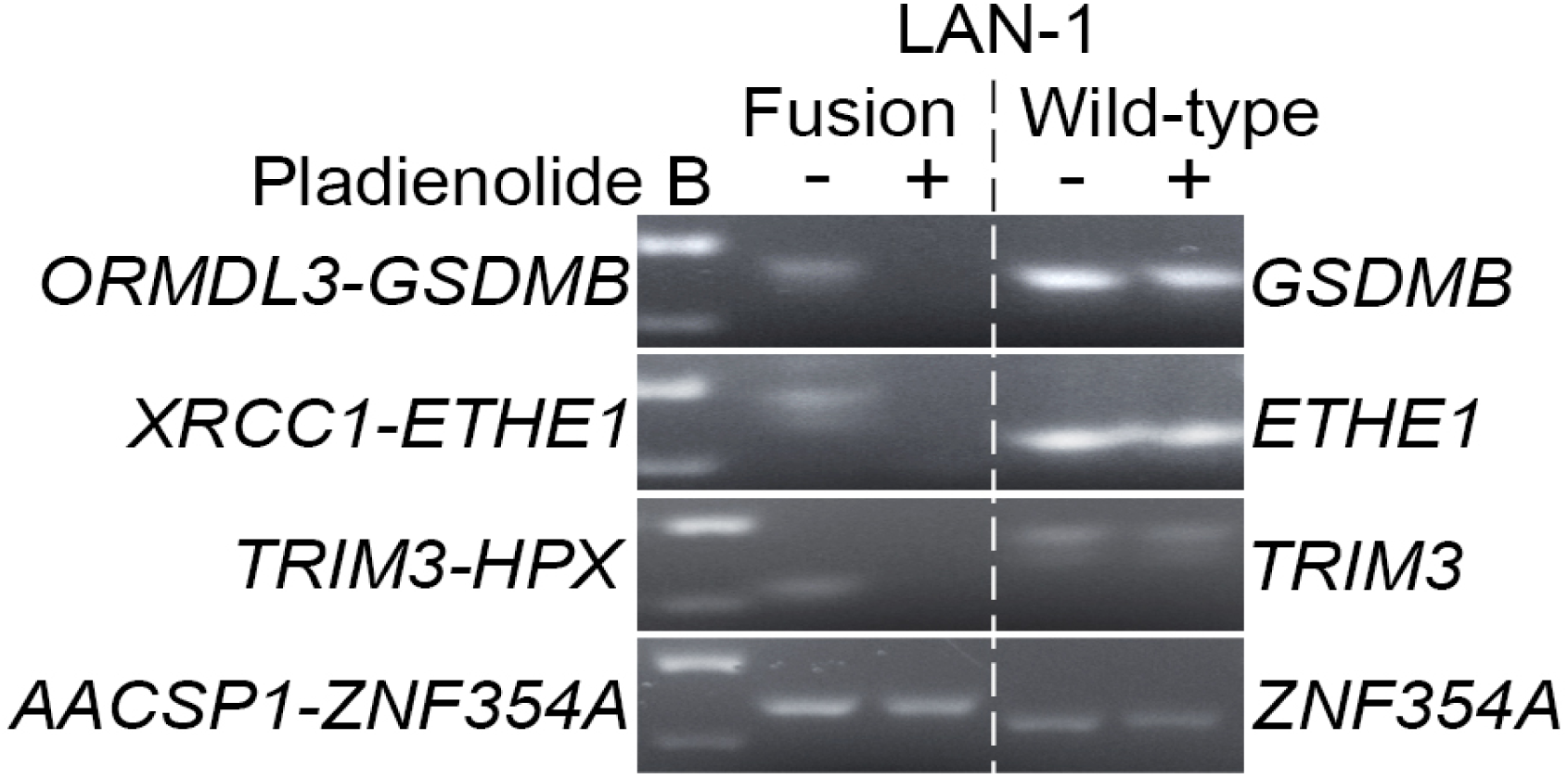
**Pharmaceutical inhibition of spliceosome activity reduces generation of fusion transcripts in LAN-1 neuroblastoma cells.** Splicing-dependent generation of fusion transcripts in LAN-1 neuroblastoma cells. LAN1 cells were treated with splicing inhibitor Pladienolide B at 5-50 nM (LAN-1) for 6 hours; RNA was isolated from harvested cells and reversely transcribed to cDNA; RT-PCR were performed with primers spanning the fusion junctions (fusion transcript) or exon-exon boundary (wild-type cognate).

---

**Table S1:** Sample information from neuroblastoma patient samples for 172 pair-end RNA-seq data from NCI TARGET project.

**Table S2:** List of fusion transcripts identified by FusionCatcher in each sample from the NB172 dataset.

**Table S3:** Frequency of fusions identified by FusionCatcher in NB172 dataset.

**Table S4:** Unique fusions in low/intermediate risk patients identified by FusionCatcher in NB172 dataset.

**Table S5:** Unique fusions in high risk patients identified by FusionCatcher in NB172 dataset.

**Table S6:** List of fusions, which encompass genes previously reported to be associated with neuroblastoma pathogenesis, identified by FusionCatcher in NB172 dataset.

**Table S7:** Tumor sample information for Validation-NB cohort.

**Table S8:** Frequency of fusions identified by FusionCatcher in Validation-NB dataset comprised of 14 neuroblastoma patient samples and 8 neuroblastoma cell lines.

**Table S9:** Frequency of fusions identified by FusionCatcher in 161 human normal adrenal glands.

**Table S10:** List of chimeric transcripts containing *TP53* as one of the fusion partners in the osteosarcoma cohort.

**Table S11:** Primers for RT-PCR validation of fusion transcripts and for cloning full length ZNF451- BAG2 as well as BAG2.

**Table S12:** Comparison of FusionCatcher detection and sequenced PCR product.

**Table S13:** Mutations identified in spliceosome associated genes in neuroblastoma (*2, 3, 35, 36*).

